# Approach direction prior to landing explains patterns of colour learning

**DOI:** 10.1101/381210

**Authors:** Keri V. Langridge, Claudia Wilke, Olena Riabinina, Misha Vorobyev, Natalie Hempel de Ibarra

**Affiliations:** Centre for Research in Animal Behaviour, Department of Psychology, University of Exeter, EX4 4QG, UK.; Department of Psychology, University of York, YO10 5DD, UK.; Division of Neuroscience & Experimental Psychology, School of Biological Sciences, Faculty of Biology, Medicine and Health, University of Manchester, Manchester, M13 9PT, UK.; Department of Optometry and Vision Science, University of Auckland, Auckland 1142, New Zealand.

**Keywords:** Insect vision, visual learning, bees, colour vision, flower patterns

## Abstract

Gaze direction is closely coupled with body movement in insects and other animals. If movement patterns interfere with the acquisition of visual information, insects can actively adjust them to seek relevant cues. Alternatively, where multiple visual cues are available, an insect’s movements may influence how it perceives a scene. We show that the way a foraging bumblebee approaches a floral pattern could determine what it learns about the pattern. When trained to vertical bicoloured patterns, bumblebees consistently approached from below centre in order to land in the centre of the target where the reward was located. In subsequent tests, the bees preferred the colour of the lower half of the pattern that they predominantly faced during the approach and landing sequence. A predicted change of learning outcomes occurred when the contrast line was moved up or down off-centre: learned preferences again reflected relative frontal exposure to each colour during the approach, independent of the overall ratio of colours. This mechanism may underpin learning strategies in both simple and complex visual discriminations, highlighting that morphology and action patterns determines how animals solve sensory learning tasks. The deterministic effect of movement on visual learning may have substantially influenced the evolution of floral signals, particularly where plants depend on fine-scaled movements of pollinators on flowers.

## 1. Introduction

Eyes and associated neural architectures have evolved in diverse ways to provide animals with adaptive views of the world. Viewing conditions are also shaped by gross morphology, since the structure of the head and body will dictate the extent of coupling between vision and movement (Land, 1999). Where the eyes and/or head are highly mobile, as in many terrestrial vertebrates, viewing direction and thus acquisition of visual information can be uncoupled from body movement - the animal can look all around while, for example, moving forward.

In contrast, other animals cannot shift their gaze independently from the body due to morphological constraints, meaning that the direction of gaze is more closely tied with their actions. This coupling is particularly pronounced in many insects which are capable of very limited head movements and lack the ability to move their eyes and lenses within the head. Although flying insects, like bees and flies, display fast and minute head movements during flight to stabilize gaze (Boeddeker et al., 2010; Boeddeker and Hemmi, 2010; Boeddeker et al., 2015; Hateren and Schilstra, 1999; Riabinina et al., 2014; Schilstra and van Hateren, 1998), to vary their viewing direction and field of view to detect image features they must change the orientation and position of the entire body. Thus, the gross movement of the insect dominates its experience of the visual environment. Insects can actively alter movement patterns to change viewpoints and actively acquire visual information (e.g. Egelhaaf et al., 2012; Land and Collett, 1997; Lehrer and Srinivasan, 1994), but this has consequences for the behavioural task at hand, and could therefore incur costs to an extent that will determine when and how active vision strategies are employed. Here we address a hitherto unexplored scenario where the efficient execution of action sequences takes a high priority in a behavioural task, asking whether that it will significantly influence an insect’s perception of readily visible objects.

Motor performance can be costly, even though it may often appear to the human observer that animals move effortlessly. For instance, air is a very viscous medium for flying insects and they have to obey the laws of aerodynamics, hence insect flight and landing manoeuvres are complicated and require a number of well-coordinated actions (Dickinson et al., 2000; Fry et al., 2005; Vance et al., 2014). For example, bees landing on a horizontal or vertical surface exhibit a sequence of highly stereotyped visually-controlled movements in order to alight successfully (Baird et al., 2013; Reber et al., 2016; Srinivasan et al., 2000). We argue that, when visual cues are not limiting, efficient motor patterns define the viewing conditions and incidentally determine what visual information is acquired for solving various behavioural tasks, such as learning the colours and patterns of a food source, a task of particular importance for bees foraging on flowers.

Colour and pattern perception in bees has been widely investigated, and it has been often assumed, implicitly and sometimes explicitly, that bees will adopt movement patterns that optimally support solving the perceptual learning tasks, e.g. approach a target in the most convenient way to view all available visual features (e.g. Avarguès-Weber et al., 2012; Dyer et al., 2005; Dyer et al., 2008; Giurfa et al., 1999a; Lehrer, 1998; Lehrer, 1999; Menzel and Lieke, 1983; Thivierge et al., 2002; Wehner, 1972; Wu et al., 2013; Zhang et al., 2004). However, the viewing conditions of individual bees will be wholly dependent upon their flight behaviour during approach and landing on the stimulus (Giurfa et al., 1999b; Hertz, 1935; Wehner and Flatt, 1977). Previous evidence suggests that bees might generally prefer to approach vertically-presented stimuli from below (Anderson, 1977; Giger, 1996), which could significantly affect perception and learning processes. This question is highly relevant for understanding the bee’s natural foraging behaviour as many flowers are tilted or vertically oriented. We show that the outcome of a learning task with flower-like colour patterns is indeed determined by the bees’ approach directions during training.

## 2. Material and Methods

### (a) Setup and training

Experiments took place indoors between October 2011 and March 2013, using bumblebees (*Bombus terrestris* Linnaeus 1758) from six colonies supplied by Koppert UK Ltd. Bees were housed in a system of Plexiglas boxes and tunnels, one of which (the ‘flight tunnel’) led to the experimental flight cage (mesh netting on a Dexion frame: 100×75×75cm). The flight cage was predominantly lit by natural daylight from a large window wall, but in addition the lab’s high-frequency lighting and three 36W diffused strip lights above the flight cage were switched on. A video camera (Photron SA-3 or Panasonic SDR-H90) was positioned perpendicular to the vertical stand with the target to record a bee’s approach from the side over the last 10 cm during training and test trials. In some of the test trials (when the test pattern’s contrast line was rotated by 90°) the bee’ choice behaviour was recorded by a second video camera from above.

Target stimuli, coloured discs (8cm diameter), and the grey background were printed on a single sheet, centred and glued to the front of a 20 x 20cm grey plastic stand. A transparent pipette nib (4mm diameter) inserted in the centre of the stimulus was backfilled with 50% sucrose solution. Eight different training patterns were used: a plain disc (Yellow or Blue) or bicolour patterns, either with the horizontal contrast line in the centre (Blue/Yellow and Yellow/Blue) or with the contrast line below or above the centre. The position of the line was at a quarter of the disc’s diameter above the bottom or below the top of the disc (3:1 diameter ratio). When the line was near the bottom of the disc the pattern was mainly blue with a small yellow section (3B:1Y), or vice versa it was mainly yellow (3Y:1B). Equally, when the contrast line was positioned in the upper half of the disc, the resulting pattern was mainly blue (1Y:3B) or yellow (1B:3Y). Blue and yellow were chosen because they are easily learnt and discriminated by bees, as confirmed by our spectral measurements (see Hempel de Ibarra et al., 2014 for details). The colours differed from each other and from the grey background in terms of both brightness and chromatic contrast for the bee eye under the illumination conditions of the flight cage.

We recorded the bees when they were close to the target, and the visual angle subtended by the target varied from 40° to above 150° as the bees approached for landing. The bees’ speed decreased linearly with decreasing distance to the target. Whilst the bees tend to pitch the abdomen much more frequently, the head’s pitch appears to be kept very steady relative to the horizontal flight direction towards the target (K. Langridge and N. Hempel de Ibarra, personal observations from high-speed video footage). Only in the very last moments of the landing manoeuvres when the bees tilt their body to position the legs at the target, the head is pitched upwards in synchrony with the rest of the body. We therefore excluded any data points closer than 0.5cm from the analysis.

All bees were marked and pre-trained individually over a few trials to fly to a uniform grey target positioned 60cm from the flight tunnel entrance, before switching to a new one displaying the coloured pattern on a grey background. Individuals were given two further practice flights before filming began.

Each bee was only trained with one type of training stimulus of 8 coloured targets described above. During ten rewarded consecutive trials each bee was filmed flying towards, and landing on, the training stimulus. In most cases, bees explored the upper half of the flight cage during this training, rather than immediately initiating the approach flight from the tunnel exit. All bees however would descend voluntarily at a larger distances from the target and approach it directly (Figure 1). Only one bee was ever present in the flight cage at any one time. Bees from several colonies participated in all colour treatments. All paper stimuli were replaced frequently (every few trials) to preclude the build-up of olfactory cues.

**Figure 1.**
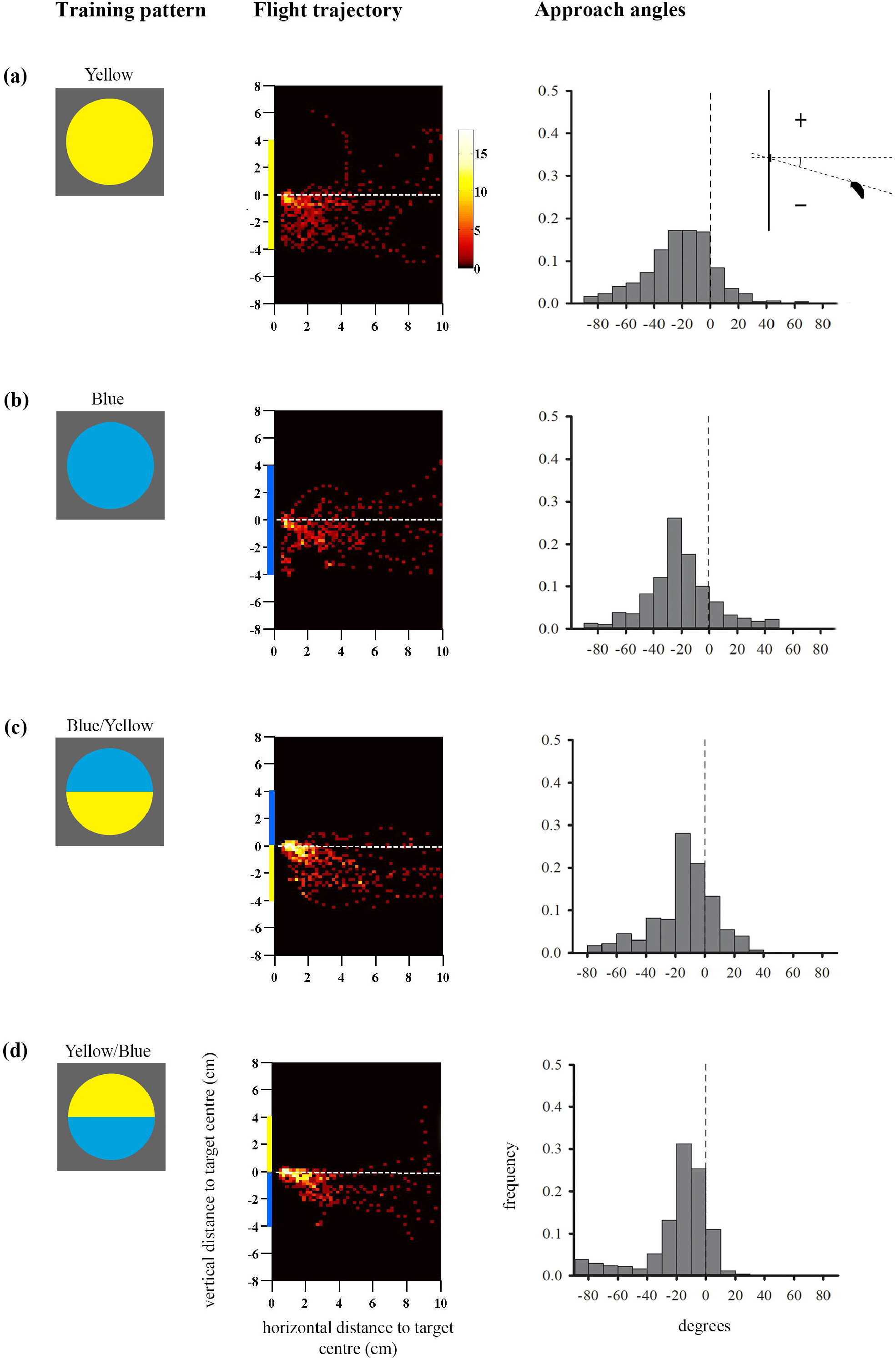
Approach flights of bumblebees trained to collect sucrose from the centre of a vertically-presented coloured disc. Training patterns were single-colour (a) Yellow or (b) Blue discs, or bicolour (c) Blue/Yellow or (d) Yellow/Blue patterns with a central contrast line, presented on a grey background. Heat-map plots depict the video capture area (filmed from the side) divided into 0.04 cm^2^ pixels denoting the frequency of positions of bees during the last training flight prior to tests (see colour scale next to Figure 1a) (n=10 bees for (a) and (b); (c) n=11 bees and (d) n=12 bees). X-axis depicts horizontal distance and Y-axis vertical distance to the target plane. Approach angles are shown on the right (see inset). Negative angles indicate a position below the contrast-line, and positive angles above the contrast line.

### (b) Colour learning tests

On completing training, each individual was subject to unrewarded tests in order to examine how the colour targets were learned. In a test the bee was presented with one of the test patterns: a single disc partitioned by either a horizontal or a vertical central boundary into two equally-sized segments, one blue and one yellow. Bees trained to single-colour stimuli had two tests with such bicolour patterns where the coloured segments were separated by a horizontal contrast line (Blue/Yellow and Yellow/Blue). Bees trained to patterns with a horizontal central contrast line were given three separate tests with bicolour test patterns that resembled the training pattern but were rotated by 180°, 90°, and 270°. Bees trained to patterns with an off-centre contrast line had two tests with bicolour test patterns with a horizontal contrast line (Blue/Yellow, Yellow/Blue) and one test with a vertical contrast line (Blue on the right, Yellow on the left). Test trials were separated by 1-3 refreshment trials with the rewarded training target, and the test sequence was varied across individuals. Test duration was three minutes from the moment of release into the flight cage. Tests with patterns where the central contrast line was horizontally oriented were filmed from the side, while test patterns with a vertical contrast line were filmed from above, such that the bees’ choices behaviour between the two coloured segments could be compared. All bees completed all training and test trials, with the exception of two individuals, which completed two tests each.

### (c) Data analysis

The approach height of the bee relative to the dorso-ventral arrangement of the colour patterns was extracted from video footage (25 fps) using a Matlab routine (see Hempel de Ibarra et al., 2009 for details), recording the position of the top line of the bee head (guided by the bee’s antennae base) in relation to the centre of the target (Figure 1).

Approach angles are defined as the angle between the head position of the bee and the contrast line within bicolour patterns, or the central horizontal line of the target for single-coloured targets, relative to the horizontal direction of approach (see inset in Figure 1a), and were analysed using Mardia-Watson-Wheeler multiple comparisons tests (with an adjusted alpha) in a circular statistics programme (Oriana V.3).

Three-minute learning tests were analysed using the behavioural data-logging freeware JWatcher (http://www.jwatcher.ucla.edu/), which calculated the total search time of the bees on the blue or yellow sector. Search time was defined as the time the bee spent flying in front of the test stimulus and visually exploring it, within 5cm distance, facing towards the pattern, during a three-minute test. Searching of the top versus the bottom colour-half was determined by the position of the top of the bees’ head relative to the contrast line. Statistical analysis of linear data was carried out using SPSS. Data met the assumptions of parametric tests unless otherwise stated.

## 3. Results

### (a) Experiment 1: single-colour and bicolour patterns with a central contrast line

Figures 1 and 2 show the approaches and test responses of bees trained to one of the single-colour and bicolour patterns where the horizontal contrast line crossed the centre of the target.

**Figure 2.**
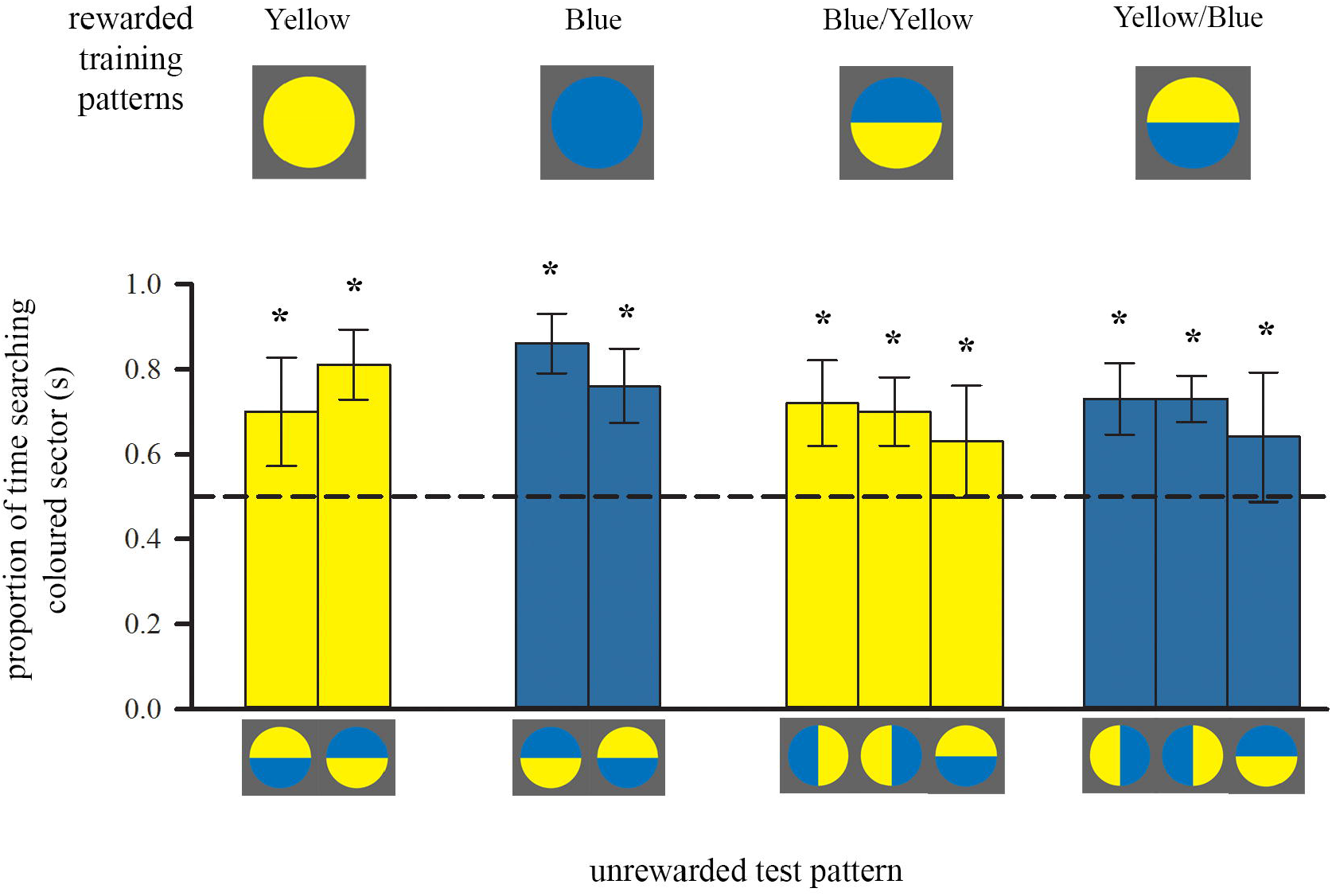
Colour preferences of trained bees in each of either two or three unrewarded tests after training to single-colour discs or bicolour patterns, respectively. Bars represent the mean proportion of time spent searching on the test pattern segment that was of either the colour of the trained single-coloured disc or the colour of the lower half of the bicolour training pattern (also depicted by bar colour), relative to the search on the other segment of the test pattern. For sample sizes see Figure 1. During each test only one of the test patterns was presented. Error bars represent ±1 standard deviation. Asterisks above bars denote a significant deviation from equal choice at α = 0.05 (Paired t-tests; see also supplementary table S1).

#### (i) Approach flight

Bees trained to collect sucrose from the centre of a vertically-presented coloured disc preferentially directed their approach towards the lower half of the pattern prior to landing (Figure 1; see also supplementary Figure S1a, b for the mean flight trajectory for each target). The flight trajectory heat maps in Figure 1 illustrate the remarkable consistency of this behaviour in bees that were not experimentally restrained in their direction of approach and landing, in response to both single-coloured discs (a, b) and bicolour patterns (c, d). The Figures strongly suggest that the approach from below-centre was inherently preferred by bees and most convenient for a successful landing.

In addition to this general trend, Figure 1 illustrates how flight paths were more streamlined and directional towards bicolour stimuli than to single-colour stimuli, presumably aided by the central contrast line. It is known that contrast lines and edges are salient visual features for bees, used to steer flight behaviour (Lehrer et al., 1990; Lehrer et al., 1985). Bees approached the lower edges of the single colour discs very closely before ascending steeply towards the centre, whereas those trained to bicolour discs began to ascend slightly earlier, generally before 2cm horizontal distance from the target, and cluster more around the target centre (Figure 1). This difference was reflected in the approach angles of the bees (Figure 1), with a significantly greater spread of steeper negative angles in response to the single-colour as compared with the bicolour targets (Blue vs. Blue/Yellow: W = 79.6, p < 0.0001; Yellow vs. Blue/Yellow: W = 35.4, p < 0.0001). There was no significant difference between bicolour pattern treatments (Blue/Yellow vs. Yellow/Blue, adjusted α = 0.0125: W = 8.7, p = 0.013).

The approach flights differed to some extent between the single-coloured blue and yellow discs (Blue vs. Yellow: W = 8.8, p = 0.012): bees exhibited a peak approach angle in response to the blue target, which was steeper than the peak angle evoked by bicolour targets (between −20° and −30°) and suggests that they approached from lower down and likely attended to the lower circle edge.

#### (ii) Colour learning tests

After ten training trials, we asked the bees what they had learned about the colour patterns by recording their search behaviour on unrewarded bicoloured discs, presenting several rotations to control for a spatial bias. Bees trained with single colours showed a significant preference for their training colour versus a novel colour (Figure 2, left; see also supplementary table S1).

Bees trained with a bicolour pattern did not search equally on both colours during tests (Figure 2, right), even though their training patterns presented equal amounts of both yellow and blue. Instead, both groups of bees showed a significant preference for the colour of the lower half of their respective training pattern: bees trained with Blue/Yellow spent significantly more time searching the yellow sectors of the test patterns for the reward, regardless of spatial position, and *vice versa* bees trained with the Yellow/Blue pattern searched more at the blue sectors (Figure 2). Colour preferences were not quite as strong as for bees trained with single-colour stimuli (F_1,39_ = 8.69, p = 0.005), suggesting that bees might also have perceived the top colour of the training pattern during approach flights. Importantly, there was a positive correlation between the amount of time individuals spent flying below the centre of the bicolour patterns over ten training flights, and the strength of their preference for the colour of the lower half (*r_s_* = 0.407, n = 23, p = 0.027).

### (b) Experiment 2: manipulation of the position of the contrast line in bicolour patterns

To investigate the causal relationship between approach flight and visual learning, we trained bees to one of four bicolour patterns with an unequal ratio of the two colours, where the contrast line was shifted into either the upper or lower half of the disc (Figures 3 and 4). We predicted that, if bees continued to approach these patterns from below centre during training, they would subsequently prefer either one or both colours presented in the lower half of the training pattern, irrespective of the overall colour distribution.

**Figure 3.**
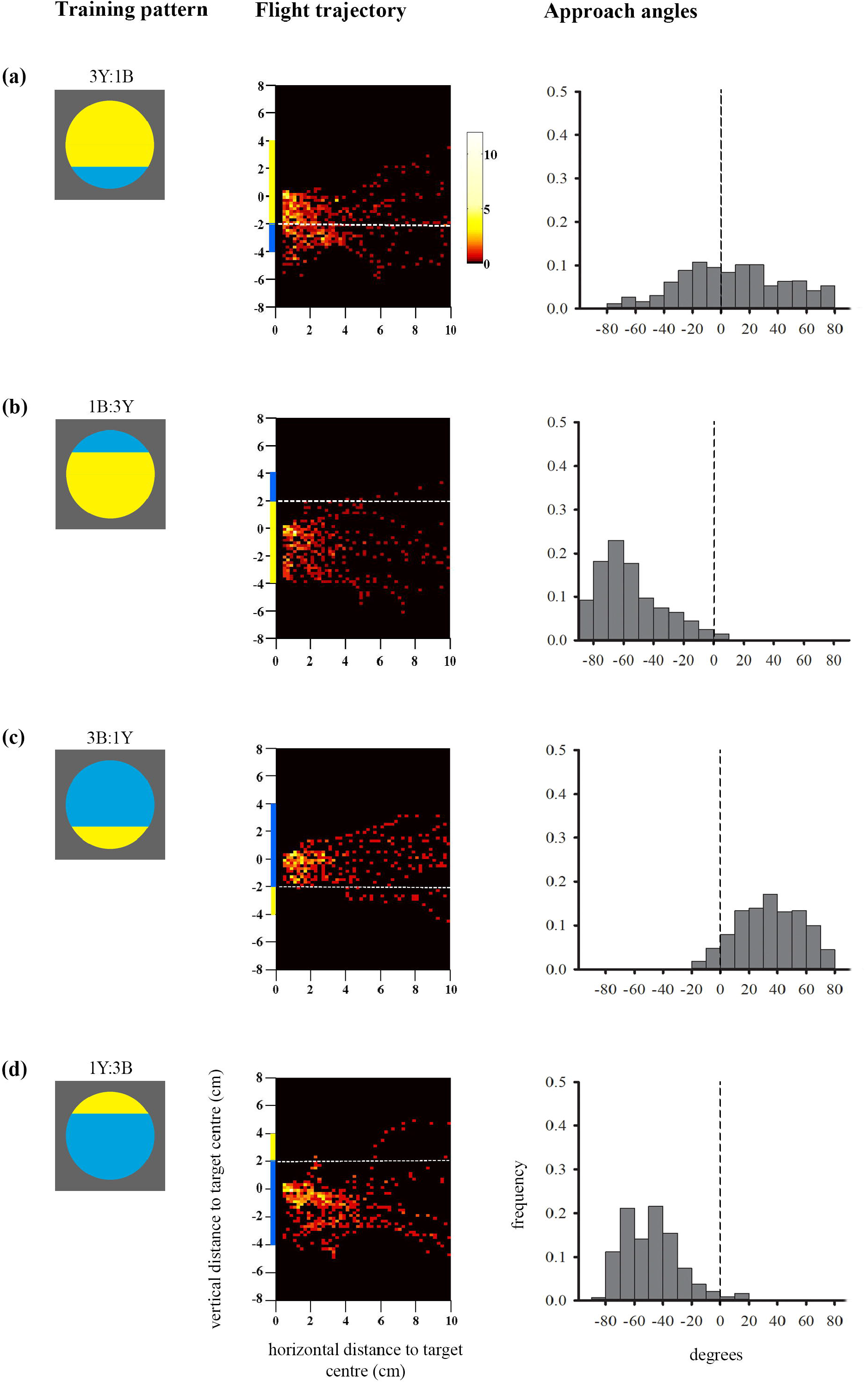
Colour preferences of bees trained with bicolour patterns with unequal segments (3:1 diameter ratio) in subsequent unrewarded tests. Bars represent the mean proportion of time spent searching on the test pattern segment that displayed the same colour as the larger segment of the training stimulus (also depicted by bar colour) relative to the search on the other segment, in each of the three tests. For sample sizes see Figure 4. Error bars represent ±1 standard deviation. Asterisks above bars denote a significant deviation from equal choice at α = 0.05 (Paired t-tests; see supplementary table S1).

**Figure 4.**
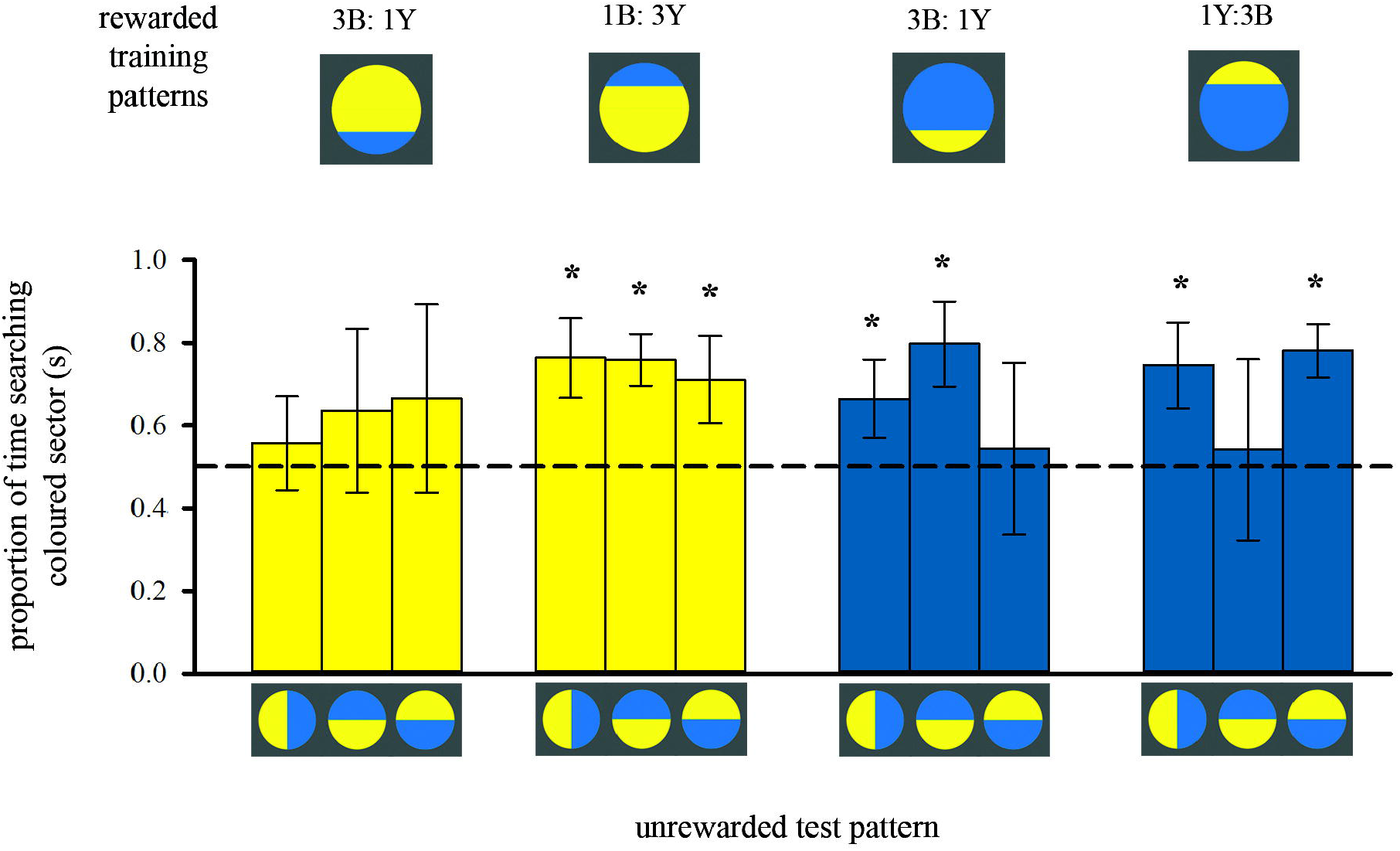
Approach flights of bees trained to collect sucrose from the centre of bicoloured patterns with an off-centre contrast line. Training patterns were (a) 3Y:1B (mostly-yellow) pattern with the contrast line and a blue segment in the bottom half or (b) 1B:3Y in the top half of the target, or (c) 3B:1Y (mostly-blue) pattern with the contrast line and yellow segment in the bottom half or (d) 1Y:3B in the top half of the target. As in Figure 1, the heat maps depict the position of the bees during the last training flight before tests (see colour scale next to Figure 1a) (n = 10 bees for all groups). White dotted lines indicate the position of the contrast line. X-axis depicts horizontal distance and Y-axis vertical distance to the target plane. Approach angle plots show the angle between the head position of the bee and the contrast line.

#### (i) Approach flight

The heat-maps in Figure 3 (a,b) show that bees approaching the mostly-yellow patterns spent considerable time viewing the whole lower half of the patterns, from the centre to beneath the lower edge (see also supplementary Figure S1c,d). When the blue segment was in the top half of the yellow pattern (1B:3Y) bees did not fly higher to view it frontally which explains the significant yellow preference shown in the subsequent tests (Figure 4). When the blue segment was present in the lower half of the pattern (3Y:1B) it was viewed equally with the yellow area below the centre of the target enabling the bees to associate both colours with the reward.

In contrast, bees trained to both mostly-blue patterns flew significantly higher, closer to the height of the target centre which contained the reward (F_1, 36_ = 8.6, p = 0.006; supplementary Figure S1d). This was particularly evident with the 3B:1Y pattern, where bees never flew below the contrast line within the last 4cm distance of the target (Figure 3c). When the contrast line was in the upper half of the pattern (1Y:3B), bees largely approached from below centre and did not fly directly in front of the yellow segment (Figure 3d), although they appeared to be more oriented towards the centre of the disc than those trained with the reversed colour arrangement in the pattern (Figure 3b).

#### (ii) Colour learning tests

Training to mostly-yellow colour patterns resulted in the predicted colour preferences (Figure 4 left; see also supplementary table S1). When the contrast line was in the lower half of the training pattern with the smaller bottom blue (3Y:1B), bees showed no significant preference for yellow in subsequent tests with bicoloured/rotated stimuli, even though yellow was the majority colour in the training pattern. Conversely, when the contrast line was in the upper half of the training pattern (1B:3Y), bees showed a significant preference for yellow in subsequent tests.

Bees trained to mostly-blue colour patterns deviated from the predicted pattern of colour preferences (Figure 4, right, supplementary table S1). In two out of three tests individuals had a significant preference for blue regardless of contrast line position during training, suggesting that they had learnt to associate blue more strongly with the reward. But the test pattern where colour positions were reversed, creating an ‘inconsistent’ spatial arrangement, elicited equal colour preferences. This result implies that for the mostly-blue patterns, bees predominantly viewed the blue area in front of them, but they also learnt the spatial distribution of the two colours around the contrast line. Indeed, variation in approach flight behaviour can explain this difference in learning performance when training bees to either mostly-blue or mostly-yellow patterns (Figure 4).

Flight patterns can explain the preference for blue in two out of the three tests for bees trained with mostly-blue patterns. However, the same bees showed equal preferences for both colours when tested with ‘inconsistent’ spatial arrangement of colours around the contrast line (Figure 4). It appears that the bees’ search behaviour in these tests was influenced not only by colour, but also by the spatial cues in the trained pattern. This is consistent with the our finding that during training to mostly-blue patterns the bees approached the contrast line differently than when trained to mostly-yellow patterns.

The approach angles differed significantly. For the mostly-blue pattern with the contrast line in the lower half (3B:1Y) there was less variation in approach angles than in the corresponding colour reversal (Figure 4a,c) (3B:1Y versus 3Y:1B: W = 151.8, p < 0.0001). Bees seem to have utilised the contrast line to guide their approach towards the central reward by positioning themselves above it, and therefore learned the spatial distribution of the colours. However, they did not fly directly in front of the small yellow segment, and therefore did not associate it as strongly with the reward as the blue colour. Furthermore, when the contrast line was in the top half of the pattern (Figure 3b,d) bees trained with the mostly-blue pattern (3B:1Y) did not fly as low and approach as steeply as those trained with the colour-reversal (1Y:3B versus 1B:3Y: W = 38.5, p < 0.0001), suggesting that they may also have attended the contrast line to guide the approach from below.

In addition to the contrast line, the outer lines of the target seem to have influenced the bees’ approach and landing manouvres. When measured with a spectrophotometer and modelled for the bee eye (see Methods), yellow was found to have a higher L-receptor (brightness) contrast for bees than blue against the grey background. The mostly-yellow patterns provided salient edge cues that differed from the small but conspicuous yellow segment in the mostly-blue patterns that were positioned either above or below the reward location. Interestingly, in the previous experiment we observed a difference in spread of approach angles between the blue and yellow single-coloured discs (Figure 1a, b) that are likely to be a consequence of the outer coloured edges against the grey background.

We conclude that the observed variations in learning performance between colour treatments are a consequence of variations in flight behaviour prior to landing. Bees modified their approach height in response to the distribution of salient pattern cues (contrast line and and salient outer edges) edges), within narrow limits dictated by their preferred landing position, and learned the colours they were incidentally exposed to as a result of this trajectory.

## 4. Discussion

We addressed novel aspects of the interaction between movement and visual perception in bees, finding that flight manoeuvres during approach and landing on a vertical target influences what they learn whilst performing a foraging task at a fully visible pattern. It has been extensively studied how walking and flying insects perform visually-guided flight control and navigation tasks. Visual information is used to avoid collisions, negotiate narrow gaps, land on a surface, or locate invisible nest or foraging sites (reviewed by Egelhaaf et al., 2012; Srinivasan and Zhang, 2004). In navigation, routes will be determined by the availability of suitable visual cues (Collett, 2010; Collett et al., 1992; Collett et al., 2006; Collett and Zeil, 1998; Harris et al., 2005; Srinivasan, 2011; Wystrach and Graham, 2012). Central-place foragers (bees and wasps) can facilitate the acquisition of visual landmark and optic flow cues required for large-scale navigation by adopting specific motor patterns (reviewed by Collett and Zeil, 1998; Land and Collett, 1997; Wystrach and Graham, 2012), and to a lesser extent, can flexibly adjust their flight behaviour for solving spatial orientation tasks (Lehrer, 1996; Lehrer and Srinivasan, 1994). Thus for active vision, for the necessary acquisition or use of specific visual cues, bees in some instances can modify the motor output, whereas the mechanism we describe here demonstrates how a necessary or efficient motor action can incidentally determine visual input. This deterministic effect of movement, particularly when bees learn about features of a rewarding target, provides a simple mechanism for explaining performance.

An intriguing example is the well-documented bias in bees to perform visual tasks better if stimuli are presented in the lower versus the upper part of the frontal visual field. This ‘dorso-ventral asymmetry’ (Giurfa et al., 1999a; Lehrer, 1999; Menzel and Lieke, 1983; Wehner, 1972) has been attributed to adaptations in central neural mechanisms for flower detection and recognition in the lower half of the bee eye, as there are no peripheral visual specialisations that could explain it. This hypothesis assumes implicitly that the dorsal and ventral visual fields of the bee are always aligned with the upper and lower halves of a target offering reward in its centre. However, the viewing conditions of individual bees will be wholly dependent upon their flight behaviour during approach and landing on the stimulus, and previous evidence suggested that bees might generally approach vertical stimuli from below (Anderson, 1977; Giger, 1996). Although the bee eye has a large field of view, which is useful for guiding movement in three dimensions, the frontal part of their compound eye has the highest visual acuity, which is best-suited for important visual tasks (Giger and Srinivasan, 1997; Graham and Collett, 2002; Land, 1997; Luu et al., 2011; Meyer-Rochow, 1981; Robert et al., 2018; Seidl and Kaiser, 1981). Thus, the part of a stimulus projecting onto the frontal areas of the bee eye would primarily determine what the bee learns from their views of visual scenes, patterns or objects, and this would differ if the bee approached from different angles. Our findings indicate that this simpler, alternative explanation for dorso-ventral asymmetries in pattern learning should not be easily discarded.

The remarkable consistency of below-centre approach paths suggests that this flight trajectory is economical and convenient for landing. The bee has to land on its legs, which are positioned ventrally, such that the head is aligned with the target centre where the bee consumes the reward. Just prior to that, the flying bee must pitch the body without losing balance, from the more horizontal angle sustained during forward flight to nearly vertical (Evangelista et al., 2010), while continuously reducing speed (Baird et al., 2013; Ibbotson et al., 2017; Srinivasan et al., 2000). An ascending bee coming from below centre can start doing this from some distance (in our experiments from about 2cm), and can easily accelerate and ascend if the landing has to be aborted. A descending bee coming from above centre would have to keep its body axis as horizontal as possible to reduce height and speed (Baird et al., 2013), leaving little time and space to swing the abdomen into the vertical pitch, thus risking loss of balance and failure to land. Flying straight towards the centre, pitching the body vertically whilst maintaining a straight trajectory from further away, could be more difficult, as slow flight speed and vertical posture could increase the aerodynamic drag downwards (Luu et al., 2011; Taylor et al., 2013). Bees that showed more central approaches in our experiments, for example in response to the mostly-blue patterns with a low contrast line, and even the few above-centre trajectories (see Figures 4c and 1a), generally dipped down just prior to landing, in order to position the legs and head correctly. It therefore seems that the optimal way to achieve the landing position is to approach from below guided by salient pattern cues, as we find here. It is noteworthy that even when the highly-salient small yellow segment was in the upper half of the colour pattern (1Y:3B pattern, see Figure 4d), the bees still did not approach above centre and specifically look at these cues. Instead, they flew slightly higher than those bees trained with the colour-reversal, sufficient to attend the contrast line and guide the approach towards the reward location. Thus although the bees adjusted their flight behaviour, this occurred within narrow limits, supporting the hypothesis that landing from below is easier than from above, and bees are constrained by the mechanics of this manoeuvre.

We conclude that viewing conditions are critical in determining what bees learn about visual stimuli, suggesting a simpler, alternative explanation to proposed cognitive mechanisms, such as position-weighting factors (Thivierge et al., 2002; Wehner, 1972), localised feature-extraction and expansion of the visual field (Giurfa et al., 1999a), or attentional focus (Morawetz and Spaethe, 2012). Detailed analyses of spatial behaviour may reveal that this mechanism underlies or influences performance in more complex tasks, such as recognising human faces, discriminating forest scenes, or using aesthetic sense to choose between Monet and Picasso (Dyer et al., 2005; Dyer et al., 2008; Morawetz and Spaethe, 2012; Wu et al., 2013). Morawetz and Spaethe (2012) admitted that they could not fully rule out simpler explanations, and suggested that flight behaviour might have played a role to shape the bees’ responses. A subsequent analysis of the data confirmed that bees varied the height in some of the tasks (Morawetz et al. 2014), which in line with our conclusions, could suggest that in those experiments bees reverted to simpler solutions for solving the learning task at hand.

There is evidence to suggest that the efficiency of the bees’ flight trajectories may be relevant for viewing and learning in more natural settings. For example, field observations commonly describe the strong directionality of bumblebees foraging on vertical inflorescences, starting at the bottom and moving upwards (Galen and Plowright, 1985; Haynes and Mesler, 1984; Pyke, 1978; Waddington and Heinrich, 1979). Flower orientation varies, and vertically-presented flowers on slopes tend to adaptively face down-slope, receiving more visitation as they offer convenient petal orientation for landing of bees moving preferentially upwards (Ushimaru et al., 2006). Observations on flowers also reveal that flower orientation influences the landing behaviour of pollinators (Ushimaru and Hyodo, 2005). It is beneficial for flowers to guide pollinator movement in a way that enhances pollen transfer (Ushimaru et al., 2009; Ushimaru et al., 2007), and the fine-scale nectar guides are generally thought to function once a bee lands (Dafni and Giurfa, 1999; Daumer, 1958; Free, 1970; Manning, 1956). Here we show that flowers may be able to exploit the tight connection between vision and movement throughout the different phases of the approach flight and landing sequence, where bees make foraging decisions. This mechanism may be decisive in how bees learn about and handle flowers, and thus develop flower-constant behaviour.

Insect vision is inherently unintuitive; the understanding of the constraints imposed on perceptual learning, as a direct result of the animals own morphology and action patterns, could provide fundamental, practical, and useful insights in a field that more recently emphasises ‘human-like’ cognitive processes. This principle is equally applicable to more familiar vertebrate groups, where species-specific viewing conditions determined by movement could provide a simple but overlooked mechanism to explain performance in perceptual learning tasks where the experimenter makes the implicit assumption that the animal views and therefore processes a given stimulus or scene in its entirety. However, viewing may differ across tasks. Birds have limited eye movements but a highly flexible neck, and so investigate objects in a very different way from mammals, by substantially moving the head to look with different parts of both eyes (Martin, 2007; Martin and Shaw, 2010; Stamp Dawkins, 2002). Head movement recruitment to shift gaze in mammals such as cats and primates is task-specific and can be blocked if energetically costly (Fuller, 1992; Oommen et al., 2004). In primates body posture and movement systematically contribute to large gaze shifts (McCluskey and Cullen, 2007). Furthermore, not all mammals have highly mobile eyes: mice, rabbit and guinea pig do not dissociate eye and head movement much (Oommen et al., 2004). Rats are widely used in studies of visual learning, despite their relatively poor visual acuity, and often display a spatial bias for learning the lower hemifield of visual stimuli, potentially as a result of their movement along the ground and subsequent bias toward viewing the lower part of a vertical stimulus (Lashley, 1938; Minini and Jeffery, 2006). Indeed, both rats and pigeons perform significantly better in visual discrimination tasks when the targets are presented horizontally or on the floor (Delius, 1992; Furtak et al., 2009). It is likely that the movement of the animal within an experimental apparatus or structured environment will have a significant effect on the outcome of learning tasks, and experiments should be designed with ethological and morphological considerations in mind.

## Acknowledgements

We thanks L. Goss, E. K. Nicholls and I. Lopes de Sousa for help during the experiments, T.S. Collett and M. Collett for helpful discussions, and the EPSRC instrument loan pool for the use of the high-speed camera. Financial support came from the BBSRC (BB/I009329/1).

